# Bird songs on the shelf: assessing vocal activity and output using data hidden in sound archives

**DOI:** 10.1101/202978

**Authors:** Oscar Laverde-R., Paula Caycedo-Rosales, Paulo-C. Pulgarín-R., Carlos Daniel Cadena

**Author notes:** Address correspondence to Oscar Laverde-R. Current address: Unidad de Ecología y Sistemática (UNESIS), Departamento de Biología, Facultad de Ciencias, Pontificia Universidad Javeriana, Bogotá D. C.

## Abstract

Understanding how often do animals emit communication signals is of critical importance to address a variety of research questions in behavioral ecology and sexual selection. However, information on vocal output, a central component of investment in signaling, is lacking for most species employing acoustic communication. Because this lack of information is partly due to logistical and methodological difficulties in monitoring animal signaling over time, developing new approaches to quantify vocal output is of special importance. We asked whether the number of recordings of avian vocalizations in sound archives and the times when such recordings were obtained reflect estimates of vocal output and temporal patterns of vocal activity obtained through systematic monitoring of wild bird populations in tropical forest sites. Based on a sample of 43 montane forest species, we found significant relationships between the number of recordings of species detected through continuous monitoring over several months and the number of recordings archived in sound collections, especially when accounting for the area of distribution of each species. In addition, daily activity patterns based on data collected through continuous monitoring over several days did not differ from those based on recordings archived in sound collections in 12 of 15 species of lowland forest birds. Annual patterns in vocal activity of two species estimated based on recordings in collections closely resembled previously published patterns. We conclude that recordings in sound collections contain valuable yet previously unappreciated information about the vocal output and temporal patterns in vocal activity of birds. This opens the possibility of using sound collections to assess vocal output and to consider it as a variable of interest in studies on the ecology and evolution of birds and other animals that use acoustic signals for communication. We encourage field workers to keep the ears wide open, and the recorders wide ready to record.

## INTRODUCTION

Understanding how often do animals emit communication signals over various time frames is of critical importance to address a variety of research questions in behavioral and evolutionary biology. In particular, birdsong is a model system for the study of animal communication and sexual selection (Read & Weary 1992; Slater 2003). Mate choice by females in many bird species is influenced by male singing behavior and song structure, with different attributes of songs being targets of sexual selection (Cardoso & Hu 2011). For example, because traits such as song length, consistency, rate, repertoire size, syllable variety, and trill syntax are attributes commonly selected by females, they are often the focus of behavioral and evolutionary studies on acoustic signals (Arvidsson & Neergaard 1991; Podos 1997; Gil & Slater 2000; Gil & Gahr 2002; Ballentine 2004; Botero *et al.* 2009; Cardoso & Hu 2011; Woodgate *et al.* 2012; Derryberry *et al.* 2012). Because singing is time-consuming, energetically costly, and may entail other costs (e.g., increased predation), studies on acoustic communication and sexual selection would benefit from understanding the costs that birds accrue when singing over longer time frames (e.g., complete breeding seasons, Krams 2001; Gil and Gahr 2002; Shutler 2011). Although some properties of single vocalizations are related to song elaboration or complexity and may thus represent partially adequate proxies of some of the costs involved in singing, they cannot fully capture the variation existing among individuals and species in singing strategies and their associated costs. What is more costly: to emit a highly elaborate song sporadically, or to emit a simple song constantly? One cannot begin to address questions like this one without basic information about when and how often do birds sing.

Patterns of circadian variation in the vocal activity of birds are not well-known, except for the fact that, generally, birds sing more at dawn (and sometimes at dusk) than at other times of the day (Aide *et al.* 2013). Daily patterns in the vocal activity of tropical birds have not been studied in detail, except for a few studies finding that canopy birds begin singing earlier than understory birds during dawn choruses (Berg *et al.* 2006), or that vocal activity in understory birds declines markedly one hour after sunrise while vocal activity in canopy birds tends to increase 1-2 hours after dawn and then declines (Blake 1992). Nonetheless, some tropical birds are more active at different times of the day (e.g., at around noon in some woodcreepers, Antunes 2008). Beyond these general observations, however, quantitative data on daily patterns in avian vocal activity are notoriously scarce, especially for tropical species.

It is generally assumed that most birds from the temperate zone exhibit marked seasonal patterns of vocal activity over the year, but this is often thought not to be the case for species from tropical, more stable environments (Stutchbury & Morton 2001). However, many tropical species show breeding seasonality in response to pulses in resource abundance and climate (Wikelski et al. 2000), and because reproduction correlates with increased vocal activity, one should expect seasonal variation in singing behavior. Some species sing throughout the year (e.g., Koloff & Mennill, 2012; Topp & Mennill, 2008), whereas others show markedly seasonal patterns of vocal activity correlated with reproduction (e.g., Negret et al. 2015; Stutchbury & Morton 2001). Even in species that sing constantly during the year, vocal activity may vary seasonally, increasing during the breeding period (e.g., Koloff & Mennill 2012; Chiver et al. 2015).

In sum, it appears clear that the singing behavior of birds needs to be described in more detail, especially in the tropics. Because this would require direct study of the behavior of individual species over long periods of time (minimally one year) and in several places, which would be highly time-consuming and rather expensive, developing alternative means to study temporal patterns in vocal activity and vocal output (i.e., how often do birds sing) would be highly desirable. Based on the idea that information associated with museum or herbarium specimens has been used to characterize the annual cycles of organisms (e.g., flowering or fruiting phenology in plants; Borchert 1996; Boulter et al. 2006; Zalamea et al. 2011), here we explore the possibility of using information archived in sound collections to describe patterns of variation in avian vocal activity and vocal output.

Sound archives are collections of field recordings of animal vocalizations focusing mainly on birds, frogs, fishes, mammals and insects. For instance, the Colección de Sonidos Animales (CSA) of the Instituto Alexander von Humboldt, Colombia, created 20 years ago, holds around 20,000 bird recordings from this country. The Macaulay Library of Natural Sounds (MLS) at Cornell University holds around 175,000 audio recordings of more than 7,500 bird species from around the world. Xeno-canto (XC; http://www.xeno-canto.org/), a website where field researchers and bird watchers share recordings of sounds of wild birds from all over the world, has accumulated ca. 315,000 recordings of more than 9,600 bird species since it was launched in May 2005. Recordings deposited in these and other sound collections have primarily been used for species identification, general bioacoustic analyses, and as records for distributional and biogeographical studies; we propose that these archives might also offer ample yet unexplored information about vocal output and temporal patterns of vocal activity.

Because recordists rely on the vocal activity of birds to record songs and calls, one may expect the number of recordings archived in sound collections to reflect how often do birds sing. For instance, the Gray-breasted Wood-Wren (*Henicorhina leucophrys*) is a common Neotropical bird that sings throughout the year; accordingly, sound collections contain a large number of recordings of this species (380 in CSA, 182 in MLS and 288 in XC as of July 2016; Table S1). The Spangled Cotinga (*Cotinga cayana*), in contrast, is a widespread species that apparently does not use acoustic signals as its main communication channel; not a single recording of it is to be found in CSA, MLS or XC. If this pattern is consistent (i.e. that species investing more heavily in vocal communication are better represented in sound archives), then one may potentially use the number of recordings in sound collections as an index of vocal output. In addition, because recordists rely on temporal patterns of vocal activity to record birds, then one may further expect that information on the date and time when recordings in sound collections were obtained would give insights about annual and daily patterns of vocal activity. However, to make valid inferences about vocal output and temporal patterns in vocal activity based on archived recordings, one would first need to validate this approach with data on vocal activity measured directly in the field.

We asked whether the number of recordings in sound archives and when such recordings were obtained reflect estimates of vocal output and temporal patterns of vocal activity obtained through systematic monitoring of wild bird populations. Specifically, we evaluated the hypothesis that one can obtain adequate estimates of vocal output and temporal patterns in vocal activity from sound archives by testing the predictions that (1) there is a positive relation between the number of recordings of each species deposited in archives and the number of vocalizations detected via direct field monitoring in three contrasting tropical forest sites, and (2) that there are no differences in the temporal distributions of recordings available in sound archives and of recordings obtained through continuous monitoring. Our results suggest that sound recordings in collections may in fact be used to assess vocal output and to describe temporal patterns of variation in the singing activity of birds, and potentially of other animals, for various types of studies.

## MATERIAL AND METHODS

We evaluated whether recordings in sound archives reflect vocal output and temporal patterns in vocal activity described on the basis of systematic monitoring of bird populations using sound recordings obtained with autonomous recording units (ARUs, Rempel et al. 2013). To accomplish this, we first used ARUs (Wildlife Acoustics SongMeter II) to (1) measure vocal output over a period of seven months (including dry and wet seasons) in a tropical montane forest and (2) to quantify daily patterns in vocal activity in two tropical lowland humid forests. We then compared our data from these field sites with information extracted from sound archives.

### Measuring vocal output

We studied vocal output of avian species over several months in a tropical montane forest in Chingaza National Park, eastern Andes of Colombia. Chingaza is located ca. 40 km east of the city of Bogotá in the departments of Cundinamarca and Meta. Approximately 190 bird species in 40 families occur in the park (Vargas & Pedraza 2004). The region has an average annual precipitation of 1800 mm, with two distinct peaks. The dry season extends from November to March with minimum rainfall in January and February, and the wet season extends from April to October with maximum rainfall in June and July (Vargas & Pedraza 2004).

We sampled avian vocal activity in the Palacio sector of Chingaza National Park (4°41’ N, 73°50’ W) in six different locations (Table 1). ARUs were located in forests between 2950 and 3170 m elevation and placed more than 500 m away from each other to avoid recording the same individuals in more than one unit. Each ARU was programmed to record for three minutes every 30 minutes. We sampled vocal activity over several months in 2013 (Table 1): February (four ARUs), April (three ARUs), May (four ARUs), July (two ARUs), and August (two ARUs). We listened to recordings obtained from 5:30 am to 12:00 m (vocal activity decreasead markedly later in the day) and identified vocalizing species (90% of avian vocalizations detected were identified to species). For each identified sound we registered the species, type of vocalization (song or call), date, and time of day. We included in analyses 43 species with more than two recordings (i.e, we excluded nine species recorded only once or twice).

**Table 1.** Number of recordings of avian vocalizations obtained through continuous monitoring by month and elevation in Chingaza National Park from February to August 2013.

| ARU<br>No | February | April | May | July | August | Total | Geographical<br>Coordinates | Elevation (m) |
| --- | --- | --- | --- | --- | --- | --- | --- | --- |
| 1 | 248 |  |  |  |  | 248 | 4°41'47"; 73°50'53" | 2950 |
| 2 | 121 | 138 | 92 |  |  | 351 | 4°41'54"; 73°50'50" | 2960 |
| 3 | 158 |  |  |  |  | 158 | 4°42'08"; 73°50'50" | 3015 |
| 4 | 233 |  | 121 | 246 | 246 | 846 | 4°41'44"; 73°50'39" | 3030 |
| 5 |  | 88 | 38 |  |  | 127 | 4°41'27"; 73°50'24" | 3015 |
| 6 |  | 131 | 132 | 150 | 50 | 465 | 4°42'42"; 73°50'21" | 3170 |
| **Total** | 760 | 427 | 383 | 396 | 296 | 2192 |  |  |

We set to test whether the number of recordings of a given species deposited in sound archives is a valid proxy of its vocal output (i.e., how often does it sing). Thus, for the 43 species detected in field recordings in Chingaza and included in the analyses, we tallied the number of recordings archived in three different sound archives: CSA, MLS and XC. CSA holds about 20,000 recordings from Colombia, with special focus in the Andes, the Amazon Basin and the Caribbean Region. MLS is one of the largest collections of bird recordings in the world, with material from the Neotropics mainly from Peru (ca. 13,000), Brazil (ca. 10,000), Venezuela (ca. 8,500), and Ecuador (ca. 6,000); Colombia is not as well represented in this collection, with only ca. 1000 recordings. XC is a web-based, rapidly growing archive with ca. 120,000 recordings from South America; in the region, the best represented countries in terms of number of recordings are Brazil (ca. 25,000), Colombia (ca. 13,000), Peru (ca. 12,000), and Ecuador (ca. 9,000).

Distributional range size likely affects the number of recordings of each species in archives (i.e., one expects more available recordings of more widespread species). Therefore, we controlled for the effect of area of the distributional range (see below) on number of recordings for each species using Nature Serve range maps (Ridgely *et al.* 2003). Because the abundance of birds may also affect the number of recordings in archives, we also sought to correct for abundance using bird-count data collected over ten months in Finca Cárpatos, Chingaza (2800-3100 m; 4°42′ N, 73°51′ W), a location very close to our study site in Palacio (Stiles and Roselli 1998; see below).

### Quantifying temporal activity patterns

We studied vocal activity along the day for avian species assemblages in two lowland forest sites. First, Barbacoas (6°41′ N, 74°21′ W) is a humid lowland forest located in Antioquia Department, in the middle Magdalena Valley of Colombia at 300 m elevation. Pastures for cattle dominate the area, but some fragments of primary forest remain. Approximately 250 bird species in 53 families have been recorded in the area (O. Laverde-R. et al., unpubl. data). The region has an average annual precipitation of 2200 mm, with two distinct peaks. The dry season extends from December to March, with minimum rainfall in January, and the wet season extends from April to October with maximum rainfall in September and October (IDEAM, http://bart.ideam.gov.co/cliciu/barran/precipitacion.htm). Second, Bahía Málaga (3°58′ N, 77°19′ W) is located on the Colombian Pacific coast in the Chocó biogeographic region, in Chocó Department. There are no published avian inventories of the area, but more than 300 species are expected to occur in the region (Hilty & Brown 1986). The region has an average annual precipitation of 7300 mm, with two distinct peaks. The region is wet year-round, but rainfall is lower from January to February, and the wet season extends from March to November with maximum rainfall in September and October (IDEAM, http://bart.ideam.gov.co/cliciu/buena/precipitacion.htm).

To sample avian vocal activity during the day, we set two ARUs in Barbacoas for 19 days (18 December 2012 to 5 January 2013) and two ARUs in Bahía Málaga for five days (2 July 2014 to 6 July 2014). ARUs were placed in tall primary forest and were programmed to record for 3 minutes every 30 minutes, from 05:30 h to 18:00 h. We listened to recordings and identified vocalizing species; for each identified sound we registered the species, type of vocalization (song or call), and time of day. We selected the 15 most frequently recorded species (i.e., those with at least 30 recordings in ARUs) for analyses of variation in vocal activity along the day (Table 2). For the above 15 lowland species, we extracted information on the time of day associated with all the recordings deposited in XC. We analyzed all archived recordings of these species (i.e., not exclusively those from our study region or from Colombia) to ensure sufficient sample sizes were available for analyses.

**Table 2.** Number of recordings obtained via continuous monitoring using ARUs in Barbacoas and Bahía Málaga and number of recordings found in the xeno-canto (XC) archive of lowland species used to study daily patterns of vocal activity.

| Family | Species | ARU | XC | Total |
| --- | --- | --- | --- | --- |
| Tinamidae | *Tinamus major* | 31 | 55 | 86 |
| Cracidae | *Ortalis columbiana* | 60 | 29 | 89 |
| Columbidae | *Patagioenas cayennensis* | 116 | 41 | 157 |
| Columbidae | *Patagioenas speciosa* | 102 | 83 | 185 |
| Ramphastidae | *Ramphastos ambiguus* | 95 | 107 | 202 |
| Thamnophilidae | *Thamnophilus atrinucha* | 69 | 94 | 163 |
| Thamnophilidae | *Myrmeciza exsul* | 74 | 238 | 312 |
| Thamnophilidae | *Myrmeciza berlepschi* | 48 | 21 | 69 |
| Tyrannidae | *Mionectes oleagineus* | 48 | 68 | 116 |
| Tyrannidae | *Todirostrum nigriceps* | 63 | 16 | 79 |
| Tyrannidae | *Tolmomyias flaviventris* | 165 | 177 | 342 |
| Tyrannidae | *Attila spadiceus* | 33 | 325 | 358 |
| Pipridae | *Lepidothrix coronata* | 83 | 63 | 146 |
| Pipridae | *Manacus manacus* | 95 | 118 | 213 |
| Trogloditidae | *Pheugopedius fasciatoventris* | 62 | 39 | 101 |
| **Total** |  | 1144 | 1474 | 2618 |

We also explored whether annual patterns of vocal activity may also be studied using recordings deposited in sound collections. We selected two species known to exhibit seasonal patterns of vocal activity along the year and which are well represented in sound collections: the Clay-colored Thrush (*Turdus grayi*) and the Great Tinamou (*Tinamus major*). The Clay-colored Thrush exhibits a marked peak in vocal activity in the first part of the year (March-April; Figure 6.1 in Stutchbury & Morton 2001), whereas the vocal activity of tinamous is known to vary seasonally, with peaks especially in the dry season (Lancaster 1964; Negret et al. 2015). Because Great Tinamou has a broad latitudinal distribution and precipitation regimes (hence likely breeding and vocal activity) differ between hemispheres, we analyzed data only from the Northern Hemisphere. We obtained the dates of recordings for these two species in MLS and XC, grouped them by month, and examined whether annual patterns of activity estimated using the recordings in archives matched expected patterns given published data.

### Statistical analyses

To evaluate the hypothesis that the information archived in sound collections can be used to estimate vocal output and temporal patterns in avian vocal activity we tested the predictions that (1) vocal output measured via continuous monitoring using ARUs correlates positively with the number of recordings in collections and (2) that the hourly distribution of recordings obtained via continuous monitoring does not differ from the hourly distribution of recordings available in collections. To evaluate our first prediction, we regressed the number of recordings in sound collections against the area of the geographic range of each species, and then regressed the residuals of this analysis on the total number of recordings counted in our field recordings. To evaluate the effect of abundance in the number of recordings in sound collections, we regressed the number of recordings in sound collections against the abundance of each species (Stiles & Rosselli 1992), and then regressed the residuals of this analysis on the total number of recordings counted in our field recordings. Variables were log-transformed to meet the normality assumptions of linear models. We also ran phylogenetic generalized least-square models (PGLS) to account for phylogenetic effects using the caper package for R (Orme *et al.* 2012) based on a comprehensive avian phylogeny (Jetz *et al.* 2012).

To examine our second prediction, we used circular statistics. First, we evaluated whether the data obtained from ARUs and from XS were uniformly distributed along the day using a non-parametric Rayleigh test of uniformity; second, we used the Watson-Williams test to evaluate the null hypothesis that the two daily activity patterns (i.e., those obtained from ARUs and XC) are not statistically different (Kovach 2011). We also used Rayleigh tests of uniformity to assess whether vocalizations of Clay-colored Robin and Great Tinamou are uniformly distributed along the year, and qualtiatively compared the annual patterns for these species revealed by recordings in MLS and XC with published data.

## RESULTS

### Vocal output

We examined 607 three-minute recordings from our tropical montane forest site in Chingaza; in 237 of these, no avian vocalizations were detected. We detected a total of 2192 vocalizations in 370 recordings obtained using ARUs (Table S1). Of this total, we identified 1522 vocalizations as songs (70%) and 433 as calls (20%); we were unable to identify 237 vocalizations (10%). Among the 1522 identified songs we detected 52 species from 19 families; for subsequent analyses we focused on 43 species having more than three recordings of songs during our sampling period (total 1512 individual songs; Table S1). For the species detected in ARUs, we found a total of 1363 recordings of songs in CSA, 1892 in MLS, and 1651 in XC.

The number of recordings of each species recorded in ARUs and in each of the sound collections were significantly and positively related (Figs. 1a-c): CSA (p<0.0001, β=0.63, R^2^=0.32), MLS (p=0.040, β=0.27, R^2^= 0.076), XC (p<0.0001, β=0.42, R^2^=0.36). Analyses correcting for size of the distributional range of species based on residuals (Figure 1) also revealed significant relationships and the explanatory power of models was considerably larger than that of models based on raw data (Figs. 1d-f): CSA (p<0.0001, R^2^=0.41), MLS (p<0.0001, R^2^=0.42) and XC (p<0.0001, R^2^=0.40). Similar results were obtained in PGLS analyses: CSA (p<0.0001, β=0.69, R^2^=0.36), MLS (p=0.002, β=0.36, R^2^= 0.14), XC (p<0.0001, β=0.48, R^2^=0.43), with the explanatory power of models increasing when we included area of distributional range as a covariate: CSA (p < 0.0001, R^2^=0.43), MLS (p<0.0001, R^2^=0.48) and XC (p<0.0001, R^2^=0.45). Analyses correcting for abundance based on residuals did not improve the explanatory power of models CSA (p = 0.0003, R^2^=0.25), MLS (p = 0.13, R^2^=0.03) and XC (p=0.0001, R^2^=0.29) even when accounting for phylogeny; CSA (p=0.07, R^2^=0.04), MLS (p=0.81, R^2^= −0.02), XC (p=0.0001, R^2^=0.23).

**Figure 1.**
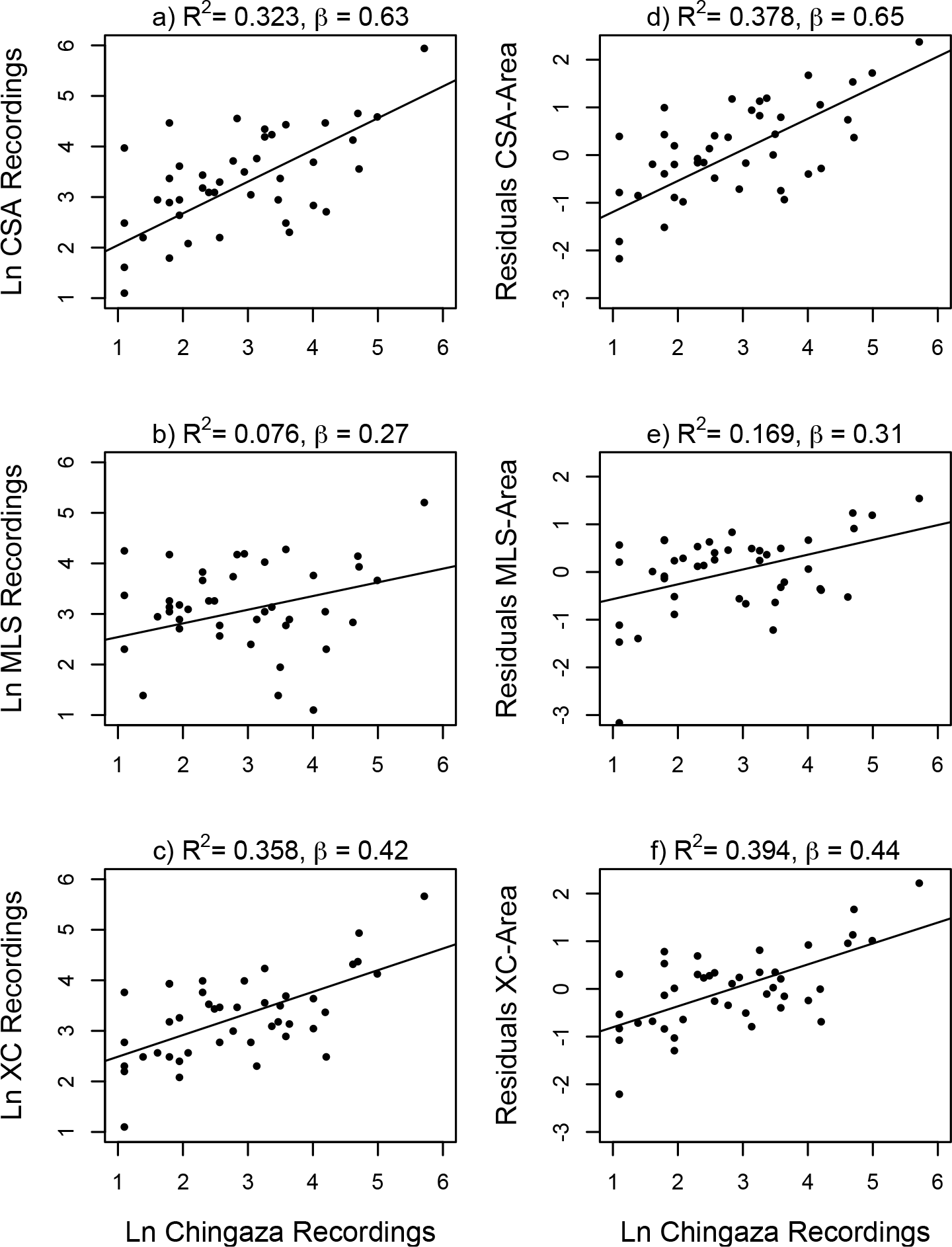
Relationships between vocal output estimated through continuous monitoring based on recordings obtained in Chingaza and the number of recordings archived in sound collections (1a – 1c), and the residual number of recordings in sound collections unaccounted for by area of distributional range (1d – 1f). Results of linear regression analyses are significant in all cases, but note the increased fit of models when accounting for area of distributional range. These patterns suggest that information in sound archives is an appropriate proxy of vocal output.

### Vocal activity

We examined 708 and 697 three-minute recordings from Barbacoas and Bahía Málaga, respectively. Of these, 431 and 430 recordings, respectively, contained avian vocalizations, resulting in a total of 1144 individual detections of species. Vocalizations recorded in the field using ARUs and those available in XC were not uniformly distributed over the day in any of our study species (Table S2; Figure 2). In 12 out of 15 species, the hourly distribution of vocalizations did not differ between data collected using ARUs and recordings available in XC, suggesting that both sources reveal similar circadian patterns of activity (Table 3). In two species of pigeon (Pale-vented Pigeon, *Patagioenas cayennensis* and Scaled Pigeon, *P. speciosa*) and one toucan (Black-billed Toucan, *Ramphastos ambiguus*) daily activity patterns were significantly different between data sets. In the pigeons, our recordings obtained using ARUs showed two clear peaks in vocal activity: one in the early morning and a second one in the late afternoon (Figure S1). Information in the XC collection did not reveal the same pattern: the peak in the morning was also clear, but the peak in the afternoon was not as evident. ARUs detected a bimodal pattern with clear peaks in the early morning and late afternoon in the singing activity of the toucan, but XC recordings showed a single peak during the mid-morning.

**Table 3.** Results of Watson-Williams circular tests for differences between the distributions of recordings obtained via continuous monitoring and based on recordings deposited in the xeno-canto archive (see Figure 2). Except for three species (two pigeons and one toucan shown in bold), daily activity patterns obtained via continuous monitoring and based on information in xeno-canto did not differ, validating the use of information in sound archives.

| Family | Species | Watson-Williams<br>F-test |
| --- | --- | --- |
| Tinamidae | *Tinamus major* | 1.999 |
| Cracidae | *Ortalis colombiana* | 2.978 |
| Columbidae | *Patagioenas cayennensis* | **5.662 \*** |
| Columbidae | *Patagioenas speciosa* | **16.768 \*\*\*** |
| Ramphastidae | *Ramphastos ambiguus* | **16.95 \*\*\*** |
| Thamnophilidae | *Thamnophilus atrinucha* | 4.233 |
| Thamnophilidae | *Myrmeciza exsul* | 0.752 |
| Thamnophilidae | *Myrmeciza berlepshchi* | 1.029 |
| Tyrannidae | *Mionectes oleagineus* | 0.649 |
| Tyrannidae | *Todirostrum nigriceps* | 0.288 |
| Tyrannidae | *Tolmomyias flaviventris* | 0.064 |
| Tyrannidae | *Attila spadiceus* | 3.639 |
| Pipridae | *Lepidothrix coronata* | 0.72 |
| Pipridae | *Manacus manacus* | 0.48 |
| Troglodytidae | *Pheugopedius fasciatoventris* | 1.523 |

**Figure 2.**
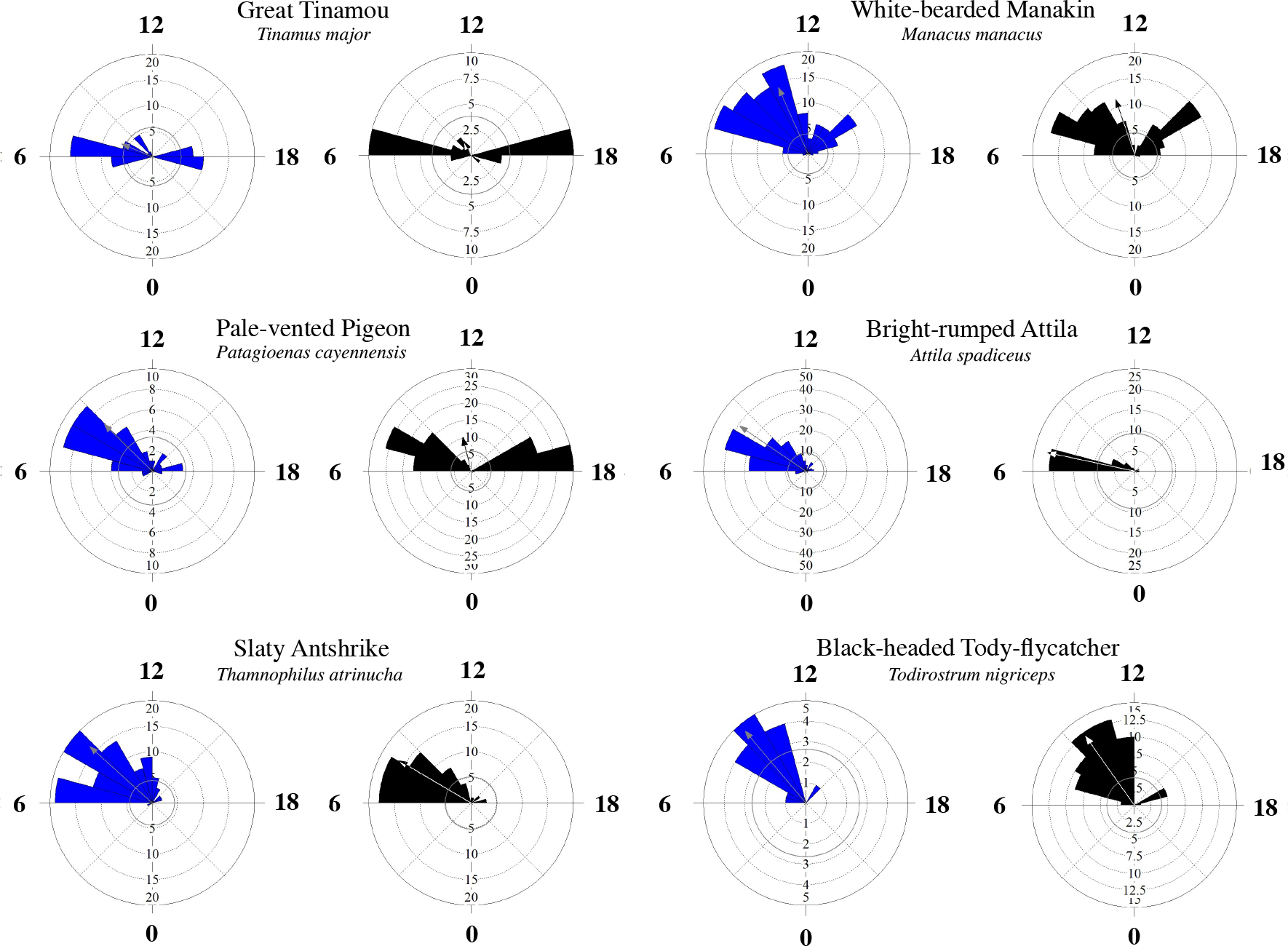
Circular plots showing daily vocal activity for six lowland species estimated via continuous monitoring (black) and based on recordings deposited in the xeno-canto archive (blue). We selected six of the 15 species studied to illustrate general patterns observed in the data. Note the strong similarity between sources of data in patterns in all species except for the Pale-vented Pigeon (*Patagioenas cayennensis*), in which the information in xeno-canto failed to detect a peak in singing activity revealed through continuous monitoring.

**Figure 3.**
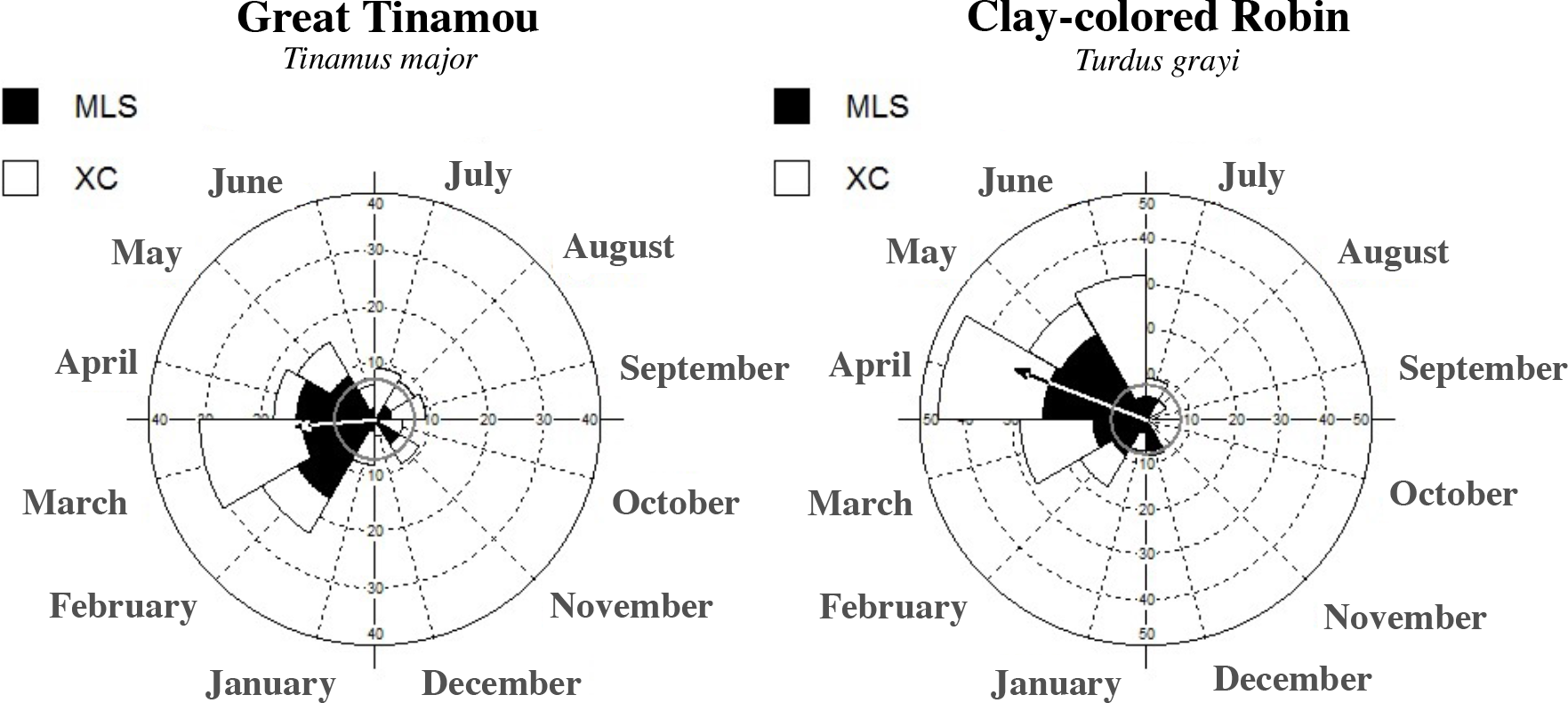
Circular plots showing annual vocal activity for the Great Tinamou and the Clay-colored Thrush in the Northern Hemisphere. Data from the MLS are shown in black and data from XC in white. Note that estimated vocal activity is very similar based on the information obtained from the two collections for both species: Great Tinamou vocal activity peaks between March and April and Clay-colored Robin vocal activity peaks in April; these patterns coincide with patterns reported in the literature based on systematic monitoring (see text).

Annual patterns of vocal activity assessed using recordings available in XC and MLS were significantly seasonal (i.e., not uniform over time) for the Great Tinamou (MLS, z= 18.816, p<0.0001; XC, z= 3.056, p=0.04) and the Clay-colored Thrush (MLS, z= 36.186, p<0.0001, XC, z= 34.877, p<0.0001). The recordings in sound archives indicate that Great Tinamou sings mostly from January to May with a peak in singing in March; the Clay-colored Thrush sings mostly from January to June with a peak in singing in April. These annual patterns based on archived recordings were similar to those described previously in the literature for species in the tinamou family (Lancaster, 1964; Negret et al. 2015) and for the Clay-colored Thrush (Stutchbury & Morton 2001).

## DISCUSSION

The energetic costs of singing and how these costs are related to song complexity or elaboration is matter of debate (Eberhardt 1994; Oberweger and Goller 2001; Ward 2004; Garamszegi et al. 2006; Hasselquist and Bensch 2008). Some studies indicate that complex songs are not necessarily more costly than simple songs (Oberweger and Goller 2001), but others suggest that complex songs can indeed be especially costly (Garamszegi et al. 2006). Regardless of any potential costs associated to song complexity, it is clear that some birds vocalize more frequently than others, implying that vocal output is a crucial variable one must consider when characterizing the variation in energy investment in vocal communication existing among species. However, vocal output is rarely evaluated in studies examining the costs of communication signals because information on how often do birds sing over various temporal scales (e.g. times of the day, months in a year) is lacking for most species. We suggest that a possible way to remedy the lack of consideration of vocal output in many studies is to use information deposited in sound collections. Our study supported the hypothesis that one can obtain meaningful estimates of vocal output and temporal patterns in vocal activity of tropical birds from information available in sound archives. This overlooked source of information may represent a crucial resource for researchers in acoustic communication, sexual selection, and other aspects of avian biology.

First, we found that recordings in sound collections can be used as a relatively accurate proxy of vocal output because the number of recordings of 43 species obtained through continuous monitoring over several months in a tropical montane forest were significantly related to the number of recordings collected non-systematically by various field workers over multiple years and deposited in sound collections; this relation was stronger when correcting the number of recordings in collections by the size of the distributional range of species but not when correcting by species abundance. Second, we found that in 12 out of 15 lowland species the similar daily patterns of vocal activity were detected using data from continuous monitoring and from sound collections. Third, for two species, circannual patterns in vocal activity determined using information in sound collections matched patterns documented in the literature based on systematic studies. Although these results encourage the use of information in sound archives to characterize vocal output and temporal patterns in vocal activity, our analyses also revealed some possible sources of bias that researchers must consider; we discuss caveats related to such biases below.

Our analyses suggest that the suitability of archived recordings to characterize vocal output may vary among collections depending on their geographic focus. For example, the relationship between recordings archived at the MLS (the collection with the lowest numbers of recordings from Colombia; ca. 1,000) and recordings we obtained through continuous monitoring was weak relative to relationships obtained using recordings archived in the XC and CSA collections, which have much larger numbers of recordings from Colombia (ca. 13,000 and 20,000, respectively). This effect may be due to regional variation in vocal activity or abundance of species in different sectors of their distribution ranges, or may reflect bias resulting from differences in recording intensity effort (i.e., collections with fewer recordings may not adequately capture patterns in vocal output due to insufficient sampling). Accordingly, the data we collected in the field for this project revealed the importance of sampling over long periods of time to adequately characterize the vocal output of species. We explored whether vocal output estimated using recordings made using ARUs over a few days reflected information in sound collections for the lowland species that we used to study daily patterns of vocal activity, finding this was not the case (F=0.04, p=0.83) even when controlling for the size of the distributional range of species (F=0.72, p=0.5). We attribute this to the fact that sound archives contain information obtained over long periods of time (i.e., multiple years including dry and wet seasons); this long-term sampling captures effects of seasonality in vocal activity that cannot possibly be revealed by data collected over short periods of time. Thus, we suggest that information in collections is not only useful to study vocal output, but also that it may provide more accurate characterizations of long-term vocal output than data collected by short-term studies.

Our work revealed that the number of recordings of species in collections is significantly related to the number of recordings of these species obtained at a single site via continuous monitoring; however, a considerable fraction of the variation in vocal output measured locally was unexplained by the frequency with which species were represented in collections. This is not unexpected given the nature of the data because collections contain information from many different sites obtained by dozens of field workers across time and space lacking the specific purpose of documenting vocal output. We explored whether aspects related to the singing behavior of species influenced the extent to which recordings in sound collections reflected vocal output estimated through continuous monitoring by examining the residuals of regression analyses relating the number of recordings in collections to vocal output assessed using ARUs, after accounting for area of distribution. First, we hypothesized that information in collections should be a worse predictor of vocal output in species with more seasonal singing activity; however, residuals did not differ between vocally seasonal and non-seasonal species (CSA, t= −0.67, p = 0.51; MLS, t = −1.10, p = 0.28; XC, t = −1.03, p = 0.30). We considered a species as non-seasonal if it was present constantly in recordings throughout the five months sampled in Chingaza, and seasonal if it was only present in recordings from 1-3 months (no species was present only in recordings from four months). Second, we also considered whether residual variation could be accounted for phylogenetic affinities; however residuals did not differ between non-passerines, suboscines and oscines (CSA, F=0.17, p = 0.83; MLS, t = −1.10, p = 0.28; XC, t = −1.03, p = 0.30).

It seemed reasonable to think that abundance influences the number of recordings in sound collections; however, we found that correcting for species abundance in our analyses did not improve the explanatory power of models relating data in sound collections to data obtained through continuous monitoring. This likely implies that birds are not recorded in proportion to their abundance by field workers contributing to sound collections; a similar conclusion has been reached for botanists collecting plant specimens for herbaria (ter Steege et al. 2011). Specifically, field workers are likely to only record few of the individuals of common species they detect, while they often try to record rare birds. That common birds may be under represented in collections while rare birds may be over represented relative to their abundance reflects not only that rarity is attractive to ornithologists, but also that most of the recordings archived in sound collections were obtained for the purpose of documenting diversity (i.e., establishing the occurrence of rare species at sites, sampling geographical variation in vocalizations of widespread species) and not for the purpose of documenting the frequency with which species were encountered.

Although daily activity patterns based on data collected through continuous monitoring were similar to those based on recordings archived in sound collections in most of our study species, this was not the case in all of them. In particular, information in collections failed to reveal a peak in singing activity in the afternoon in two species of pigeon and a toucan. Recordists, like most birds, are more active during the morning, and this certainly has an impact on the daily patterns of activity one may infer based on recordings in sound collections. Our data and recordings in sound collections reveal that tropical birds exhibit varying strategies in vocal activity through the day (Figure 2, Figure S1); some species sing in the early morning and late afternoon, others during the whole morning, and a few probably during the whole day. Therefore, to the valuable recommendations for recording bird sounds offered by previous workers (Parker III 1991; Budney and Grotke 1997), we would add the importance of recording constantly along the day and of recording as many singing birds as possible. By doing so, recordists may considerably increase the usefulness of their material for studies on various aspects of avian biology.

The idea that one may use information in sound collections to study vocal seasonality along the year as shown by our analyses focused on Great Tinamou and Clay-colored Thrush offers exciting opportunities for future study. For example, information in sound collections may allow one to assess the effect of climate on singing behavior within and across sites; because singing correlates with reproduction, information in sound collections may further be used as a proxy to examine geographic variation in breeding activity and its consequences for population differentiation (Quintero et al. 2014). In the long term, sound recordings in collections may allow one to monitor changes in vocal activity patterns and annual phenological cycles in wild bird populations in relation to local and global change.

We conclude that recordings in sound collections contain valuable information about the vocal output of birds, which opens the possibility of considering vocal output as a variable of interest in studies on the ecology and evolution of birds and other animals that use acoustic signals for communication. Sound collections also contain much information about temporal patterns in vocal activity over various time frames; this information is relevant to studies in many areas of animal biology. Just as traditional museum specimens capture valuable information about the time in which they were collected (Pyke and Ehrlich 2010), properly archived sound recordings may be considered acoustic specimens with great potential to be used in various ways in the future. Traditional specimens have been used for purposes that original collectors could not have imagined, including the documentation of the effects of environmental contaminants (Berg et al. 1966; Ratcliffe 1967) or the prediction of shifts in species distributions as a consequence of climatic change (Peterson et al. 2002). The same could be true of future studies on vocal activity and vocal output based on properly curated and publicly available sound collections. Now that sound recordings are relatively easy to collect and archive, we encourage field researchers to record animal sounds along the day and the year, and to make recordings and associated information available to the public. To finalize, we emphasize that much of data one may extract from recordings in sound collections is waiting to be used; part of the information needed for many different studies is already sitting on the shelf.

## ACKNOWLEDGMENTS

We thank Chingaza National Park for allowing us to record bird songs. Greg Budney and Matt Medler of the Macaulay Library of Natural Sounds at Cornell University, and the Colección de Sonidos Ambientales of the Instituto Alexander von Humboldt gave us unrestricted access to the information archived in their collections. We thank to Ken Rosenberg of the Cornell Lab of Ornithology for the loan of some recording equipments. We thank all the recordists who have deposited their recordings in these archives and who have made them available through xeno-canto. We are grateful to Laura Céspedes, Simón Quintero, Juan Ignacio Areta, Mecky Hollzmann, Trevor Price and Alex Kirschel for asistance in the field.

